# Sequestration of LINE-1 in novel cytosolic bodies by MOV10 restricts retrotransposition

**DOI:** 10.1101/2021.11.09.467897

**Authors:** Rajika Arora, Maxime Bodak, Laura Penouty, Cindy Hackman, Constance Ciaudo

## Abstract

LINE-1 (L1) are autonomous retroelements that have retained their ability to mobilize. Mechanisms regulating L1 mobility include DNA methylation in somatic cells and the Piwi-interacting RNA pathway in the germline. During pre-implantation stages of mouse embryonic development, however, both pathways are inactivated leading to a critical window necessitating alternate means of L1 regulation. We previously reported an increase in L1 levels in *Dicer_*KO mouse embryonic stem cells (mESCs). Intriguingly this was accompanied by only a marginal increase in retrotransposition, suggestive of additional mechanisms suppressing L1 mobility. Here, we demonstrate that L1 Ribonucleoprotein complexes (L1 RNP) accumulate as aggregates in *Dicer_*KO cytoplasm along with the RNA helicase MOV10. The combined overexpression of L1 RNAs and MOV10 is sufficient to create L1 RNP aggregates in stem cells. In *Dicer*_KO mESCs, MOV10 is upregulated due to the loss of its direct regulation by miRNAs. The newly discovered post-transcriptional regulation of *Mov10* expression, and its role in preventing L1 retrotransposition by driving novel cytosolic aggregation affords alternate routes to explore for therapy and disease progression.

## Introduction

Approximately 17-20% of human and mouse genomes are composed of Long Interspersed Nucleotide Elements 1 (LINE-1 or L1) ^1,2^. These elements, ranging from 6 to 7 kb in length, encode enzymatic activities necessary for retrotransposition. In mouse, L1s are composed of a 5’ untranslated region (UTR) harboring an RNA Polymerase II (Pol II) promoter encoding a bicistronic transcript. The two open reading frames (ORF) encode for L1 ORF1 protein that is speculated to function as an RNA chaperone and L1 ORF2 protein that has endonuclease and reverse transcriptase activities. The transcript harbors a 3’UTR and a poly adenylation (poly(A)) signal. Only a full-length poly(A) transcript is capable of transposing. Upon export from the nucleus, L1 RNA is translated in the cytoplasm. L1 RNA, ORF1 and ORF2 proteins associate to form ribonucleoprotein particles (L1 RNPs), which are imported back together into the nucleus. Once in the nucleus, the L1 RNA is reverse transcribed and integrated into a new genomic location by a coupled reverse transcription. During this mobilization mechanism, the retrotransposon sequence is prone to truncations and inversions, resulting in the insertion of mutated copies unable to jump a second time ^3,4^. Nevertheless, 100 ^5^ and 3000 ^6^ full length L1 elements in human and mouse genomes, respectively, retain the ability to encode the machinery necessary for production of the RNA intermediate, its reverse transcription, and consequent integration into a new genomic location. In mouse, active L1s are divided into three subfamilies: Tf, Gf and A, which are defined by the variable sequence and numbers of monomers (tandem repeat units of 200 bp) contained in their 5’UTR ^7–9^.

While transposable elements are indispensable for genome variation and evolution, rogue and/or rampant transposition leads to disease ^3^. Elucidating mechanisms that regulate L1 transcription and mobility have been an active area of research since their discovery. DNA methylation in somatic cells and Piwi-interacting RNA (piRNA) pathway in the germline are well established regulators of L1 retrotransposition ^10–12^. At the blastocyst stage of embryonic development however, both the above mentioned pathways are inactivated leading to a window necessitating alternate mechanisms of L1 regulation. The microRNA (miRNA) effector protein DICER has been implicated in modulating expression of L1 during this stage of development ^13^. MicroRNAs are 21-24 nucleotide (nt) long Pol II transcripts that play a major role in fine-tuning gene expression post-transcriptionally ^14,15^. Briefly, miRNAs are transcribed as primary (pri) miRNAs and processed into precursor (pre) miRNAs by DGCR8/DROSHA microprocessor complex in the nucleus. Upon export into the cytoplasm DICER cleaves pre-miRNAs to give rise to mature miRNAs. The mature miRNA duplex is loaded onto ARGONAUTE (AGO) proteins, upon unwinding of the duplex, one of the two strands is degraded. Along with accessory proteins, AGO loaded with the guide miRNA strand forms the RNA-induced silencing complex (RISC) and acts as the effector. Base pairing of miRNA at its seed sequence with complementary miRNA response elements (MREs), typically found in the 3’UTR sequence of mRNAs induces translational repression or mRNA degradation. Pre-implantation mouse embryos deleted for *Dicer* present an upregulation of L1 elements ^16,17^. In human cancer cells, miR-128 was shown to regulate L1 transposition via two mechanisms. Firstly, miR-128 repressed L1 expression directly by binding to a noncanonical binding site in L1 ORF2 RNA ^18^ and secondly, miR-128 bound to a canonical binding site in the 3’UTR sequence of Tnpo1 an import factor that regulates entry of L1 RNP complex into the nucleus post translation ^19^. This mode of regulation via miR-128 however does not appear to be conserved in mESCs ^20^. Recently, the direct binding of miRNA let-7 to L1 mRNA was shown to impair L1 ORF2 translation and consequently retrotransposition ^21^. Since processing of pri-let7 miRNA to mature let-7 miRNA is blocked in mESCs ^22^, this mechanism of fine-tuning L1 expression is also not conserved in mESCs. To delve deeper into the role of *Dicer* in regulating L1 during embryonic development our laboratory utilized mouse embryonic stem cells (mESCs) as a model. In *Dicer*_Knockout (KO) mESCs, while a 6-8 fold increase in L1 transcription was observed, a concomitant increase in the rate of retrotransposition was not uncovered ^13^. In this study, we demonstrate that miRNAs are involved in the regulation of L1 retrotransposition in mESCs through the direct regulation of the RNA helicase MOV10. Upon loss of miRNAs, MOV10 is strongly upregulated and accumulates in the cytoplasm of mESCs, driving sequestration of L1 RNPs into novel aggregates, thereby preventing L1 mobility.

## Results

In order to better understand why the strong upregulation of L1 RNAs does not lead to a subsequent retrotransposition in *Dicer*_KO mESCs ^13^, we looked at the localization of L1 RNA and protein in Wild type (WT) and mutant cells. We probed for L1 RNA derived from the Tf L1 family by RNA Fluorescent in Situ Hybridization (RNA FISH) along with L1 ORF1 protein by indirect immunofluorescence (IF). While in WT mESCs, we observed diffused signal for both L1 Tf RNA and ORF1 protein, they co-localized as L1 ribonucleoprotein (L1 RNP) foci in cytoplasm of the two independent *Dicer*_KO clones (Fig.1A). The median number of L1 RNP foci in the cytoplasm per cell in *Dicer*_KO1 and *Dicer*_KO2 mESCs was 9 and 7 respectively as compared to 0 in WT cells. Additionally, in 30-35% of *Dicer*_KO clones, L1 RNP were observed to co-localize in larger foci (Fig.1A). These observations led us to hypothesize that sequestration of L1 RNP in the cytoplasm of *Dicer*_KO mESCs is preventing L1 retrotransposition.

**Figure 1.**
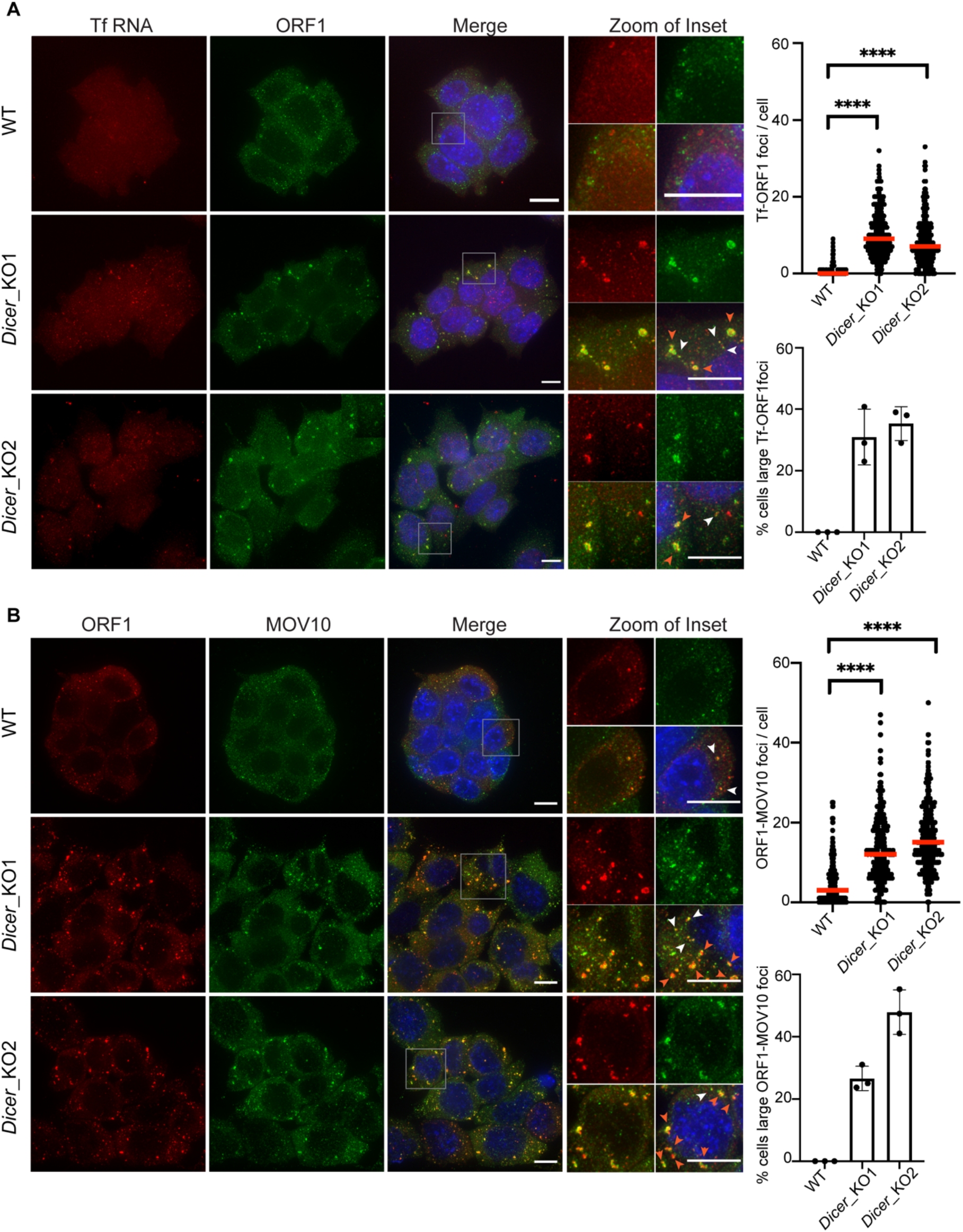
L1 RNP accumulate as cytoplasmic foci in *Dicer*_KO mESCs. **(A)** Maximum intensity projections across Z stacks of example images from indicated mESCs stained for L1 Tf RNA (red) combined with immunostaining for L1 ORF1 protein (green) and nuclei stained with DAPI (blue). The grey square marks position of the inset. White arrow heads point to cytoplasmic foci where L1 RNA and ORF1 protein co-localize. Red arrow heads point to relatively larger sized L1 RNP foci. Data collected from 275 WT, 304 *Dicer*_KO1, 311 *Dicer*_KO2 cells from three independent experiments are depicted as scatter plots where circles are single data points representing number of co-localized L1 Tf-ORF1 foci in the cytoplasm per cell, red bar is median for the distribution. P-value was determined using Mann-Whitney *U* test and **** represent p-value < 0.0001. In *Dicer*_KO cells, L1 RNA and protein co-localize in variably sized foci in the cytoplasm. Bar graphs are mean values of percentage of cells with large L1 Tf-ORF1 foci co-localizing in the cytoplasm. Dots represent data from three independent experiments, error bars are standard deviations. Scale bar 5 μm. **(B)** Maximum intensity projections across Z stacks of example images from indicated mESCs immunostained for L1 ORF1 (red), MOV10 (green) and nuclei stained with DAPI (blue). The grey square marks position of inset in the zoomed image. White arrow heads point to cytoplasmic foci where L1 ORF1 and MOV10 co-localize. Red arrow heads point to relatively larger sized L1 ORF1-MOV10 foci. Data collected from 293 WT, 295 *Dicer*_KO1, 295 *Dicer*_KO2 cells from three independent experiments are depicted as scatter plots where circles are single data points representing number of co-localized L1 ORF1-MOV10 foci in the cytoplasm per cell, red bar is median for the distribution. In *Dicer*_KO cells, L1 ORF1 and MOV10 proteins co-localize in the cytoplasm. P-value was determined using Mann-Whitney *U* test and **** represent p-value < 0.0001. Bar graphs are mean values of percentage of cells with large L1 ORF1-MOV10 foci co-localizing in the cytoplasm. Dots represent data from three independent experiments, error bars are standard deviations. Scale bar 5 μm.

To characterize L1 RNP foci we aimed to identify other cellular components that might share their location with them. We therefore tested if known interactors of human L1 proteins might colocalize with L1 RNP cytoplasmic foci in *Dicer*_KO mESCs ^23–26^. Amongst the list of candidates interacting with both L1 ORF1 and L1 ORF2 ^26^, we looked at RNA helicases UPF1 and MOV10 by IF. While UPF1 was observed to have diffused cytoplasmic staining (data not shown), MOV10 co-localized with L1 RNP in the cytoplasm of *Dicer*_KO mESCs (Fig. 1B). Further analysis revealed MOV10 to co-localize with L1 ORF1 protein in *Dicer*_KO cells with a median of 3 foci in WT cells and 12 and 15 respectively in *Dicer*_KO1 and *Dicer*_KO2 mESCs. Percentage of cells with large ORF1-MOV10 foci was 26-47% in the two *Dicer*_KO lines (Fig. 1B). The higher frequency of ORF1-MOV10 foci as compared to Tf-ORF1 foci in *Dicer*_KO cells is most likely due to the lower sensitivity for detecting Tf RNA by RNA FISH. Since MOV10 co-localization with L1 ORF1 foci in *Dicer*_KO mESCs was high and due to the absence of good antibodies available for L1 proteins raised in hosts other than rabbit for co-staining IF experiments, we further used MOV10 as a proxy for L1 RNP localization.

Localization of L1 RNP as cytoplasmic foci was previously reported for human L1 proteins upon their ectopic overexpression in HEK293T cells ^27^. L1 ORF1 foci were furthermore shown to co-localize with stress granules and RNA-binding proteins including components of the RISC complex ^27^. To assess the nature of the observed mouse L1 RNP foci, we co-stained WT and *Dicer*_KO mESCs for G3BP1, a marker for stress granules ^28^, along with MOV10. The signal for G3BP1 was mainly diffused cytoplasmic in both WT and *Dicer*_KO mESCs, indicating that unlike human cancer cells, mouse L1 ORF1-MOV10 foci are not stress granules (Fig. 2A). However, treatment with 0.5mM Sodium Arsenite for 20 minutes to induce stress caused MOV10 to co-localize with G3BP1 as cytoplasmic bodies in *Dicer*_KO cells (Fig. 2A). These data led us to hypothesize that L1 RNP foci in *Dicer*_KO mESCs might be poised but are not as yet mature stress granules.

**Figure 2.**
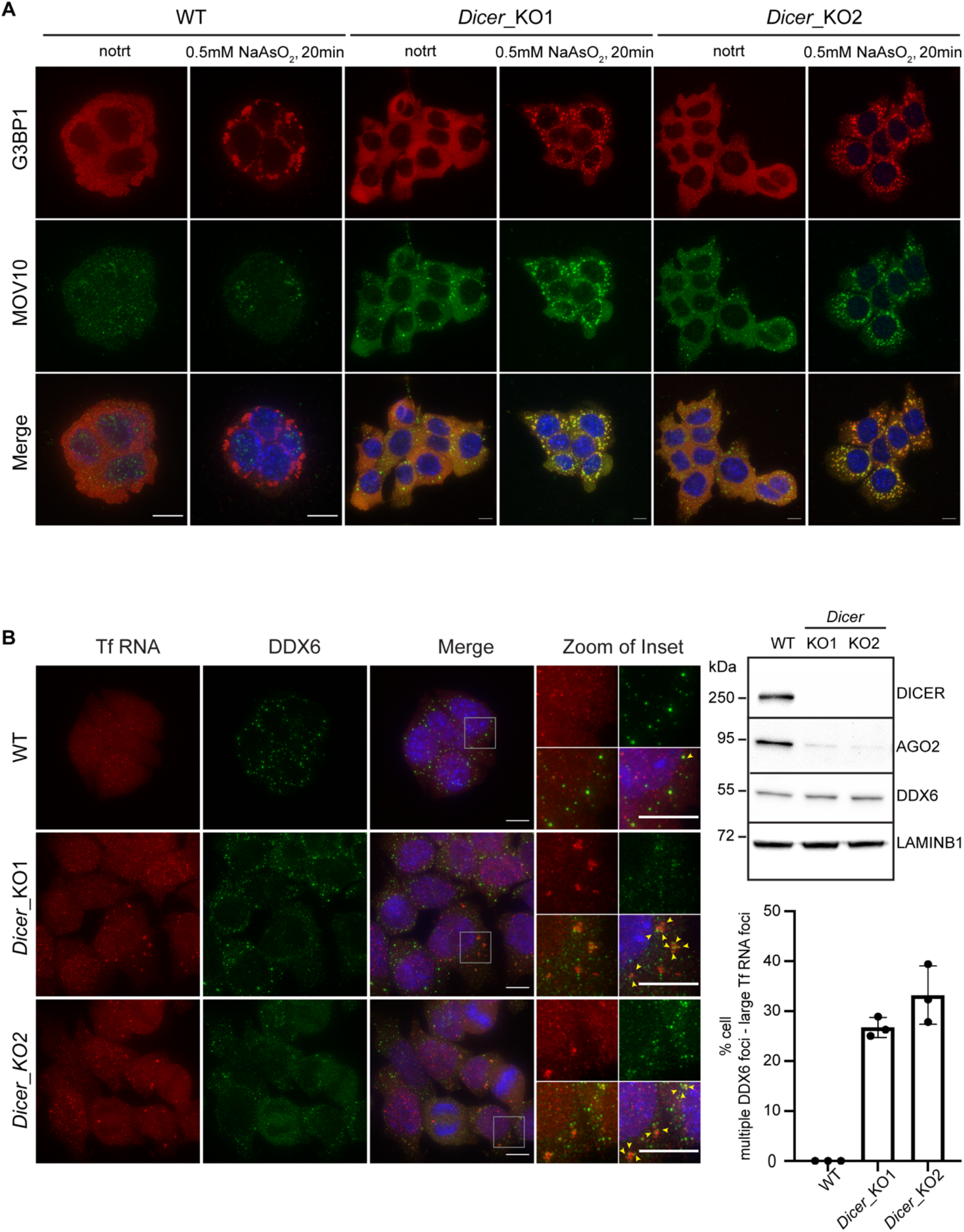
Cytosolic L1 RNP foci are poised to be stress granules that co-localize with multiple small DDX6 foci. **(A)** WT and *Dicer*_KO mESCs were treated with 0.5mM Sodium Arsenite (NaAsO2) for 20 minutes or left untreated prior to fixation with formaldehyde. Maximum intensity projections across Z stacks of example images from indicated mESCs immunostained for G3BP1 (red) and MOV10 (green) with nuclei stained with DAPI (blue). Diffused cytoplasmic staining of G3BP1 was observed in all untreated samples, while MOV10 was found to localize in cytoplasmic foci in *Dicer*_KO cells. Treatment with Sodium Arsenite resulted in co-localization of G3BP1 and MOV10 in stress granules in *Dicer*_KO cells. Images are representative of 3 independent experiments. **(B)** Representative Western Blots out of 3 independent experiments showing low AGO2 protein levels in *Dicer*_KO as compared to WT mESCs (right side). No change in protein levels for DDX6 was observed, LAMINB1 served as loading control. Immunoblotting with Anti-DICER antibody was performed to confirm the fidelity of the KO clones. On the left side, maximum intensity projections across Z stacks of example images from indicated mESCs stained for L1 Tf RNA FISH (red) combined with immunostaining for a resident protein of P-bodies, DDX6 (green) and nuclei stained with DAPI (blue). The grey square marks position of the inset. Yellow arrow heads point to cytoplasmic foci where L1 RNA and DDX6 protein co-localize. Multiple small DDX6 foci were observed to co-localize with large L1 Tf RNA foci in the cytoplasm of *Dicer*_KO but not in WT mESCs as depicted in the bar graph. Dots represent data from three independent experiments with percentage computed from at least 94-150 cells per cell line per experiment, error bars are standard deviations. Scale bar 5 μm.

Partitioning of stress granule proteins as liquid-liquid phase separation (LLPS) is emerging as a main driver for shifting dynamics from being near soluble to condensate formation thereby impacting their biological function ^29^. RNA and RNA binding proteins (RBPs) are key components of these cytoplasmic condensates ^30^. Recently, by microscopy and NMR spectroscopy, human L1 ORF1 protein was shown to form liquid droplets *in vitro* in a salt dependent manner ^31^. To test whether L1 ORF1 foci in mESCs undergo similar LLPS, we treated *Dicer*_KO mESCs with 3% 1,6 Hexanediol for 15 minutes, a concentration at which proteins undergoing LLPS have been previously observed to change solubility from being in foci to becoming diffused in mESCs ^32^. No overt change in L1 ORF1-MOV10 foci was observed in cells treated with 1,6 Hexanediol (Extended Data Fig.1A), suggesting that L1 ORF1-MOV10 foci are not LLPS condensates.

Human L1 ORF1 protein are also known to associate with Processing Body (P-body) enriched mRNAs ^33^. While elucidation of the functional relevance of P-Bodies is an active area of research, it is well established that these cytoplasmic granules also undergo LLPS ^34^. Since the L1 RNP foci are not sensitive to 1,6 Hexanediol treatment and most likely not undergoing LLPS, our data argues against L1 RNP foci being components of P-body in mutant mESCs. Additionally, the protein ARGONAUTE2 (AGO2), a known component of P-bodies ^35^ and an effector of the miRNA biogenesis pathway, is required for P-body formation ^36,37^. In *Dicer*_KO mESCs due to the absence of miRNAs, AGO2 protein levels are reduced and the protein destabilized ^13^ (Fig. 2B). However protein levels of DDX6, another known constituent of P-bodies ^38^, were unchanged as compared to WT cells (Fig. 2B). We therefore looked at the cellular localization of DDX6 to assess P-body integrity and association with L1 RNP foci. Unlike WT cells where DDX6 formed droplet like foci characteristic of P-bodies in the cytoplasm, in *Dicer*_KO cells, DDX6 was more diffusely localized in the cytoplasm. In 26-32% of *Dicer*_KO mESCs, multiple small DDX6 foci were observed co-localizing with larger L1 Tf RNA foci (Fig. 2B). The partial co-localization with DDX6 in cells with low AGO2 levels suggest that L1 RNP foci are not canonical P-bodies, corroborating earlier studies enumerating the requirement of intact miRNA biogenesis in P-body fidelity ^36,37^.

Finally, we ascertained that L1 RNP foci were not autophagosomes ^39^ as LC3B a marker for autophagosomes did not co-localize with MOV10 in mESCs by IF (Extended Data Fig. 1B). We therefore called L1 RNP present in cytoplasmic foci of *Dicer*_KO mESCs, aggregates as they contain an assembly of RNA and proteins without undergoing phase separation.

L1 upregulation is amongst the many changes in gene expression observed upon deleting *Dicer* in mESCs ^13^. To parse out whether as observed in human cultured cells overexpression of L1s was sufficient for cytoplasmic sequestration ^23,27^, we engineered WT mESCs to endogenously upregulate L1 using CRISPRa (L1^UP^) (Fig. 3A, Extended Data Fig. 2A). We designed single guide RNAs (sgRNAs) to target dCas9 fused with VP160 to the 5’UTR sequence of the L1 Tf family (Extended Data Fig. 2B). For the generation of independent clones (Cl), L1^UP^ Cl1 cells were transfected with one sgRNA, while two sgRNA pairs were used to upregulate L1 in L1^UP^ Cl2. A 2.5-fold increase in L1 Tf transcript levels as compared to the control cell line (Ctrl) transfected with an empty sgRNA vector was observed (Extended Data Fig. 2C) in L1^UP^ clones. Given the sequence homology of the three L1 families, we also observed a 3-fold increase in transcript levels of L1 A family, while the increased expression of L1 Gf family was found to be statistically significant for only Cl1 (Extended Data Fig. 2C). While L1 transcript levels in L1^UP^ cells was lower than in *Dicer*_KO (Extended Data Fig. 2C), expression of L1 ORF1 protein in L1^UP^ was similar to that observed in *Dicer*_KO cells (Fig 3A).

**Figure 3.**
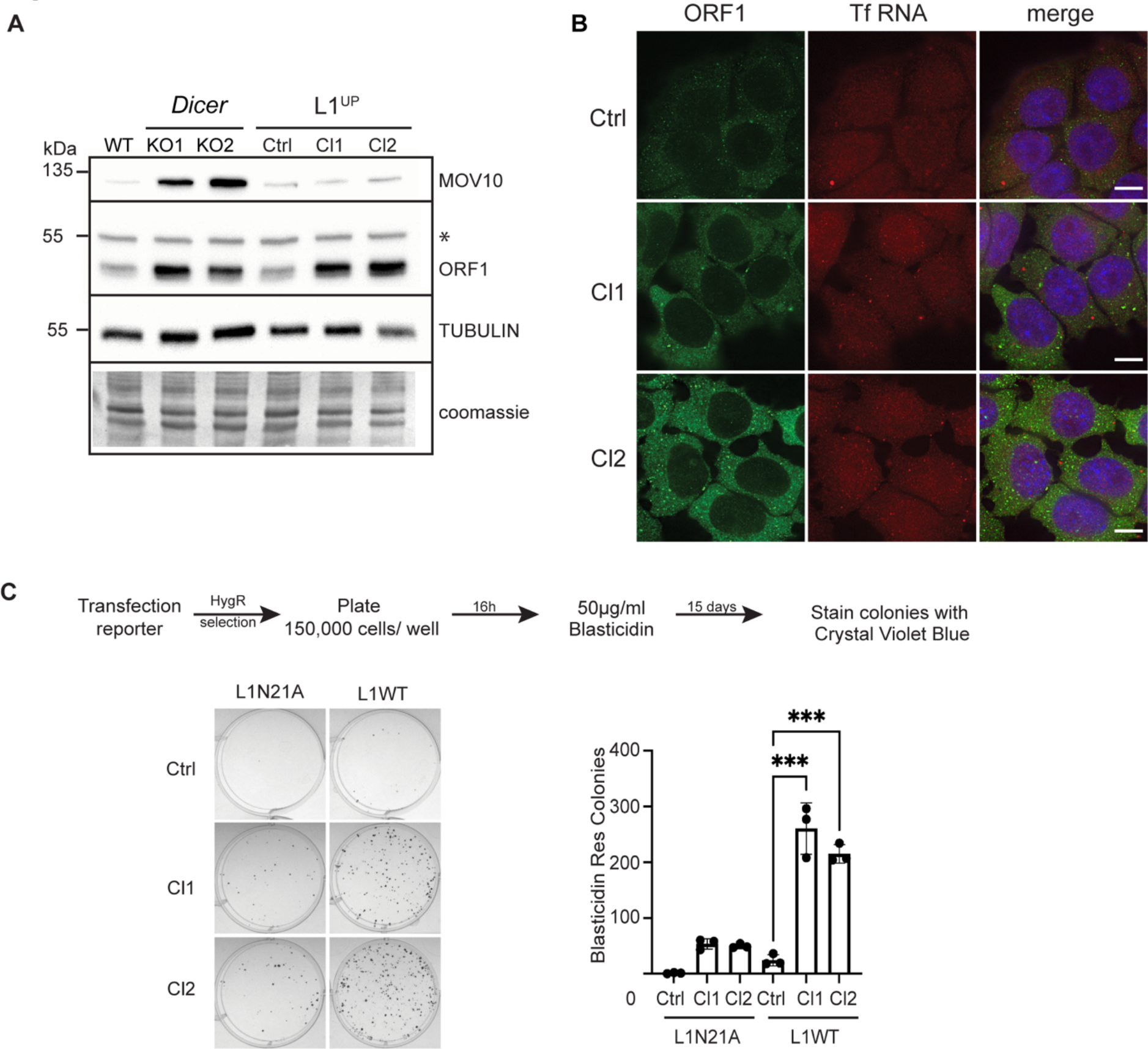
Endogenous L1 upregulation leads to L1 retrotransposition. **(A)** Representative Western Blots out of 3 independent experiments showing L1 ORF1 and MOV10 protein levels in the indicated cell lines. Immunoblot with antibody recognizing TUBULIN and coomassie stained membranes depict the loading, asterisk marks position of non-specific band in the ORF1 immunoblot. While higher L1 ORF1 levels are observed in *Dicer*_KO and L1^UP^ cells compared to WT and Ctrl cells respectively, MOV10 overexpression is only observed in *Dicer*_KO cells. **(B)** Maximum intensity projections across Z stacks of example images from indicated mESCs immunostained for L1 ORF1 protein (green) combined with RNA FISH for L1 Tf RNA (red) and nuclei stained with DAPI (blue). No aggregation of L1 RNP in the cytoplasm is observed in L1^UP^ cells. Scale bar 5 μm. **(C)** Representative images of BlastR colonies stained with crystal violet blue of indicated cell lines is shown on the left. Cells were transfected with either mutant reporter plasmid (L1N21A) or retrotransposition competent reporter (L1WT) as shown in the scheme with timeline for the experiment on the top. Bar graphs on the right depict the average number of BlastR colonies, dots are mean values obtained from 3 independent experiments, error bars are standard deviations. P-value was determined using unpaired student t-test and *** represent p-value < 0.001.

To assess if L1 elements upregulated with CRISPRa were competent for retrotransposition, we primarily performed Northern Blot analysis and observed that like in *Dicer*_KO mESCs, full length L1 transcripts were being overexpressed ^13^ (Extended Data Fig. 2D). Importantly, this level of upregulation of L1 RNA was not sufficient to cause L1 RNP accumulation in cytoplasmic aggregates in L1^UP^ mESCs (Fig. 3B). Using a plasmid based retrotransposition assay ^40^, we tested if in the absence of L1 RNP cytosolic sequestration there was an enhanced rate of L1 retrotransposition in the engineered L1^UP^ mESCs. We transfected Ctrl, L1^UP^ Cl1 and L1^UP^ Cl2 with either wild type JJ-L1SM (L1WT) or a plasmid with mutation in ORF2 rendering it incompetent for jumping (L1N21A) that carried *Blasticidin* resistance (BlastR) as a reporter gene and *Hygromycin* (HygR) as a selection marker ^41^. Unlike in *Dicer*_KO cells ^13^, L1 upregulation was accompanied by an increase in the rate of mobility depicted by the higher number of BlastR colonies observed in L1^UP^ Cl1 and Cl2 as compared to Ctrl mESCs (Fig. 3C). BlastR colonies observed in the two L1^UP^ cell lines transfected with L1N21A reporter confirm previous observation of mobilization of mutant L1s aided by endogenous full length L1s in the cell, but at relatively low frequencies ^42^. To conclude, forced endogenous upregulation of L1 active elements in WT mESCs is not sufficient to create L1 RNP cytoplasmic aggregates and leads to an increase in retrotransposition.

Given that upregulation of L1 in mESCs was not sufficient to induce L1 RNP aggregation in the cytoplasm (Fig. 3B), and our finding that MOV10 co-localized with L1 RNP in *Dicer*_KO cells (Fig. 1B), we speculated that cytosolic aggregation of L1 RNP might be driven by the upregulation of MOV10 observed in *Dicer*_KO mESCs at RNA and protein levels (Extended Data Fig. 2E, 3A). MOV10 upregulation in *Dicer*_KO mESCs was confirmed by RTqPCR analysis (Extended Data Fig. 2F). In addition, no changes in MOV10 expression were observed either at RNA (Extended Data Fig.2F) or protein levels in L1^UP^ mESCs (Fig. 3A). We therefore ruled out L1 overexpression as the driver for MOV10 upregulation and investigated the role of miRNAs in post-transcriptional regulation of *Mov10* as miRNA biogenesis is impaired in *Dicer_KO* mESCs.

Using TargetScan software ^43^, we identified multiple miRNAs (miR-138-5p, miR-30-5p, miR-16-5p and miR-153-5p) as predicted to target the 3’UTR sequence of *Mov10* (Fig. 4A). The relative expression of each miRNA in WT cells was determined using previously published small RNA sequencing data from our laboratory ^44^ (Extended Data Fig. 3A). MiR-16-5p and miR-30-5p are highly expressed in WT mESCs compared to the intermediate expression of miR-138-5p, and the low expression of miR-153-3p (Extended Data Fig. 3A). We tested whether the predicted miRNAs might directly regulate *Mov10* expression by performing a luciferase reporter assay ^45^. We subcloned the 3’UTR sequence of *Mov10* downstream of the *Renilla luciferase* reporter gene in a plasmid that also encoded *Firefly luciferase* as a normalizer. Transient transfection of this plasmid along with the respective miRNAs into HEK293T followed by measurement of the respective luminescence showed that for the tested mimics, RENILLA expression was significantly sensitive to transfection with miR-16-5p and miR-153-3p (Fig. 4B). To corroborate that the upregulation of MOV10 in *Dicer*_ KO cells is indeed mediated by miRNAs and is not a consequence of noncanonical function of *Dicer*, we tested whether a similar upregulation of MOV10 is present in *Drosha*_KO cells where the canonical miRNA biogenesis pathway is also impaired ^46^.

**Figure 4.**
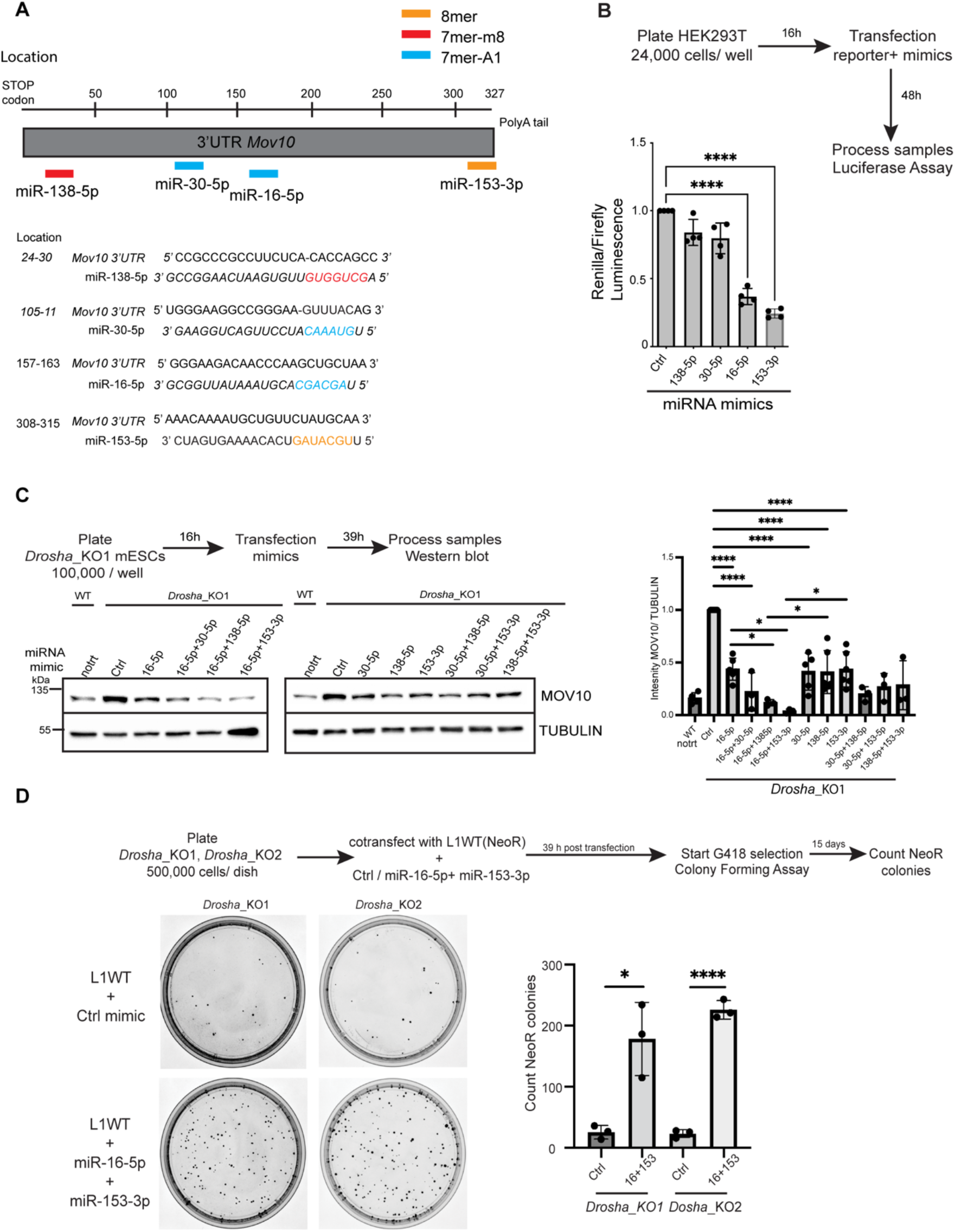
Multiple miRNAs regulate MOV10 expression and L1 retrotransposition in mESCs. **(A)** Schematic of 3’UTR sequence of *Mov10* RNA helicase. Location of miRNA response elements (MREs) for mouse miR-138-5p (red), miR-30-5p (blue) miR-16-5p (blue) and miR-153-3p (orange) predicted to target *Mov10* (ENSMUST00000168015.8) are color coded based on seed type matching for respective miRNAs. **(B)** Schematic showing design and timeline of luciferase assay performed in human Hek293T cells to assay direct miRNA mediated repression of *Mov10*. Bar graphs show the average relative luminescence of reporter gene *Renilla* to which the 3’UTR sequence of *Mov10* was fused, normalized by Firefly luminescence, where relative ratio observed for transfection with control (Ctrl) mimic was set to 1. Each dot on the bar graph is the mean from 4 independent experiments, errors are standard deviation. P-values were calculated using an unpaired t-test and **** are p-values < 0.0001. Renilla expression was sensitive to transfection with miR-16-5p and miR-153-3p. **(C)** Schematic showing design and timeline for processing samples for WB analysis in *Drosha*_KO1 cells. *Drosha*_KO1 were transfected either with Ctrl mimic or with indicated miRNA mimics either singly or in pairs. For comparison protein from untreated (notrt) WT cells was also run on the same blot. Blots were probed with anti-MOV10 and anti-TUBULIN antibodies. Bar graphs show mean intensity of MOV10 normalized by TUBULIN from 3 independent experiments relative to transfection for the Ctrl mimic that was set to 1. P-values were computed using ordinary one-way ANOVA test comparing the mean of each sample to the mean of Ctrl, and the mean of doubly transfected mimic with its singly transfected counterpart. * depict p-value < 0.05, and **** depict p-value < 0.0001. MOV10 expression was found to be sensitive to transfection with all four transfected mimics. MiR-16-5p was found to down-regulate expression of MOV10 synergistically with both miR-138-5p and miR-153-3p, to levels similar to those observed in WT mESCs. **(D)** Schematic summarizing the experiment and timeline followed for colony forming assay in *Drosha*_KO mESCs transfected with either Ctrl mimic or miRNA mimics for miR-16-5p + miR-153-3p along with L1WT plasmid bearing NeoR gene as reporter. Representative images of NeoR colonies stained with crystal violet blue of indicated cell lines are shown on the left. Bar graphs on the right depict the average number of NeoR colonies, dots are mean values obtained from 3 independent experiments, error bars are standard deviations. P-value was determined using unpaired student t-test and * represent p-value < 0.05, **** represent p-value < 0.0001. Downregulation of *Mov10* expression due to transfection with indicated miRNA mimics in *Drosha*_KO cells resulted in an increase in the rate of mobility of L1 elements.

Western blot (WB) analysis on *Drosha*_KO cells revealed that MOV10 is indeed upregulated in these cells (Extended Data Fig. 3B). Finally, to confirm miRNA mediated regulation of *Mov10* expression, we transiently transfected *Drosha*_KO mESCs with the respective miRNA mimics either singly or in pairs and measured MOV10 expression. Unlike previously observed with the luciferase assay, expression of MOV10 was downregulated upon transfection with each of the four tested miRNA mimics (Fig. 4C). Interestingly, only paired transfection of miR-16-5p with miR-138-5p or miR-153-3p acted synergistically to reduce MOV10 protein levels down to WT levels (Fig. 4C). Collectively, these data reveal a role for miRNAs in fine-tuning MOV10 expression in mESCs, explaining the observed MOV10 upregulation in *Dicer*_KO and *Drosha*_KO mESCs (Fig. 3A, Extended Data Fig. 3B).

Given the upregulation of MOV10 and L1 ORF1 in *Drosha*_KO cells as compared to WT (Extended Data Fig. 3B), we next assessed if L1 RNP correspondingly also aggregate in the cytoplasm of these miRNA mutants. We performed IF with L1 ORF1 and MOV10 antibodies in two independent *Drosha*_KO clones and observed MOV10 co-localizing with L1 RNP in the cytoplasm of *Drosha_*KO mESCs (Extended Data Fig. 3C). The median ORF1-MOV10 aggregates per cell were 21 and 12 in *Drosha*_KO1 and *Drosha*_KO2 mESCs respectively (Extended Data Fig. 3C). Percentage of cells with large ORF1-MOV10 foci was 31-44% in the two *Drosha*_KO lines (Extended Data Fig. 3C), similar to that observed in *Dicer*_KO cells (Fig. 1B).

To confirm our hypothesis that aggregation of L1 RNP driven by MOV10 overexpression was preventing L1 retrotransposition, we examined whether restoring MOV10 expression in *Drosha*_KO cells would allow L1 mobilization. We used a plasmid based retrotransposition assay ^40^ and transiently co-transfected *Drosha*_KO1 and *Drosha*_KO2 with pCEP-L1WT reporter plasmid that carried *Neomycin* resistance (NeoR) ^41^ as a reporter along with either Ctrl mimic or mimics for miR-16-5p and miR-153-3p together to downregulate MOV10 expression. 500,000 cells were plated for each condition for the colony forming assay and media was supplemented with G418 39 hours post transfection. The mean NeoR colonies obtained 15 days post selection were 25 and 23 in the two *Drosha*_KO clones transfected with Ctrl mimics from 3 independent experiments. A statistically significant increase in NeoR colonies in cells transfected with miRNA mimics was observed with the mean increasing to 178 and 226 in the two clones respectively (Fig. 4D). Our results are in line with data from human cancer cells supporting the role for *Mov10* as a negative regulator of retrotransposition ^23,47–50^, and to our knowledge, the first to report a role for miRNAs in fine-tuning *Mov10* expression.

Mature miRNAs might regulate multiple mRNAs and an mRNA can be targeted by several miRNAs ^51^. While we show that transfection with miR-16-5p and miR-153-3p mimics downregulates MOV10 expression leading to increased L1 mobility, we cannot unequivocally rule out that changes in expression of another gene targeted by these miRNAs might be responsible for the observed increase in transposition. To assess if MOV10 expression is sufficient to induce L1 RNP aggregation in the cytosol, we transiently transfected Ctrl, L1^UP^ Cl1, L1^UP^ Cl2 mESCs with a plasmid encoding HA tagged human MOV10 (HA-MOV10). In IF experiments with an antibody against HA to detect exogenously expressed HA-MOV10 along with anti-L1 ORF1 antibody, we detected HA-MOV10-ORF1 aggregates in the cytoplasm of L1^UP^ Cl1 and L1^UP^ Cl2 significantly more than in Ctrl cell line (P-val < 0.001). The median number of foci observed in Ctrl was 6 per cell while in the two L1^UP^ clones this was 15 (Fig. 5A). Additionally, the morphology of the larger HA-MOV10-ORF1 aggregates observed in L1^UP^ clones was reminiscent of those observed in *Dicer*_KO mESCs (Fig. 5A, Fig. 1B). To prove that MOV10 induced L1 RNP aggregation restricts L1 mobility, we then transiently co-transfected Ctrl, L1^UP^Cl1 and L1^UP^Cl2 mESCs with JJ-L1WT reporter plasmid that carries *BlastR reporter* ^41^ along with either Empty Vector (EV) or HA-MOV10 plasmids. The mean BlastR colonies was 35 and 29 for the two L1^UP^ clones and 2 in Ctrl cells, corroborating our earlier observation of increased L1 mobility in L1^UP^ cells as compared to Ctrl (Fig. 5B, Fig 3C). Importantly, a statistically significant decrease in BlastR colonies was observed in L1^UP^ clones transfected with HA-MOV10 when compared to EV with a mean of 1 BlastR colony obtained from the transfection in both the clones (Fig. 5B). Together, our data implicate that MOV10 is playing a direct role in cytosolic sequestration of L1 RNP thereby restricting retrotransposition and maintaining genome integrity in mESCs (Fig. 5C).

**Figure 5.**
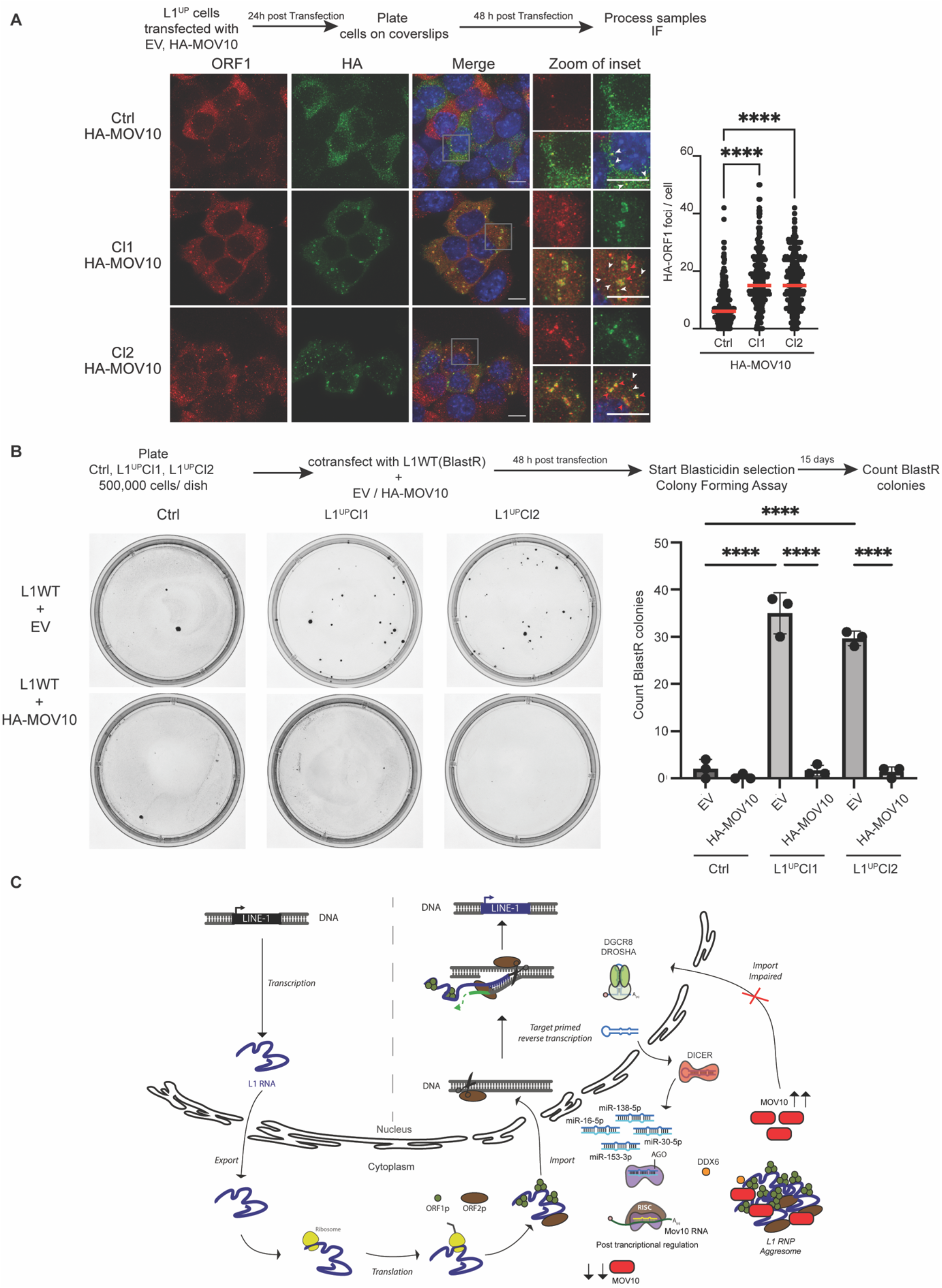
MOV10 upregulation is sufficient to create L1 RNP aggregates in L1^UP^ mESCs abrogating L1 retrotransposition. **(A)** Scheme for transfection and timeline for processing samples for IF. Maximum intensity projections across Z stacks of example images from indicated mESCs stained for L1 ORF1 (red) combined with immunostaining for HA (green) to detect ectopically expressed MOV10 tagged with HA at the N-terminus, and nuclei stained with DAPI (blue). White arrow heads point to cytoplasmic foci where L1 ORF1 and HA-MOV10 protein co-localize. Red arrow heads point to relatively larger sized HA-ORF1 foci. Data collected from 289 Ctrl, 275 L1^UP^ Cl1 and 296 L1^UP^ Cl2 mESCs from three independent experiments are depicted as scatter plots where circles are single data points representing number of co-localized HA-ORF1 foci in the cytoplasm per cell. Red bar marks median for the distribution. P-value was determined using Mann-Whitney *U* test and **** represent p-value < 0.0001. Statistically significant increase in cytoplasmic L1 ORF1 aggregates was observed upon ectopic expression of HA-MOV10 in L1^UP^ as compared to Ctrl mESCs. **(B)** Schematic summarizing the experiment and timeline followed for colony forming assay in Ctrl and L1^UP^ mESCs transfected with either Empty Vector (EV) or HA-MOV10 plasmid along with L1WT plasmid bearing BlastR as reporter. Representative images of BlastR colonies stained with crystal violet blue of indicated cell lines are shown on the left. Bar graphs on the right depict the average number of BlastR colonies, dots are mean values obtained from 3 independent experiments, error bars are standard deviations. P-value was determined using unpaired student t-test and, **** represent p-value < 0.0001. Upregulation of MOV10 in L1^UP^ mESCs restricted L1 retrotransposition. **(C)** The life cycle of L1 retrotransposition is depicted. Only full length L1 elements get transcribed driven by the promoter residing in its 5’UTR sequence. The bicistronic L1 RNA is exported from the nucleus into the cytosol and translated to give rise to L1 ORF1 (ORF1p) and L1 ORF2 (ORF2p) proteins. The L1 RNA and proteins form a complex (L1 RNP) and are imported back into the nucleus. Endonuclease activity of ORF2 nicks the target DNA and using a mechanism referred to as Target primed reverse transcription a new copy of L1 element is inserted into the genome via a copy past mechanism of mobilization ^3,12^. A key regulatory step for retrotransposition is the import of L1RNP back into the nucleus. The canonical miRNA biogenesis pathway illustrates the miRNAs discovered in this study that regulate expression of RNA helicase *Mov10* a known modulator of L1 mobility. In the absence of miRNAs when either DICER or DROSHA proteins are deleted in mESCs, both L1 and MOV10 expression are upregulated. Our data suggests that in microRNA mutant mESCs MOV10 induces L1 RNP aggregate formation in the cytoplasm, the impaired import consequently prevents L1 retrotransposition despite high L1 expression. While DDX6 was also found to co-localize with the larger L1 RNP particles, identification of molecular partners and biochemical activities intrinsic to the L1 RNP aggregates should unveil the bottle neck afforded to prevent import.

## Discussion

The role of MOV10 in inhibiting retrotransposition in human tissue culture was discovered almost ten years ago ^23^. Since then, multiple reports have corroborated this seminal function, where it participates either directly or along with protein partners in curbing retrotransposition ^47–50,52,53^. Here, we discover cytosolic-body formation induced by MOV10 as a novel line of defense for sequestration of L1 RNP particles to prevent deleterious L1 retrotransposition. It appears that L1 RNP aggregates in miRNA mutant mESCs are different from those observed upon ectopic overexpression of MOV10 and L1 in human cancer cells as the latter unlike in our study were found to be stress granules.

MOV10 is a known interactor of proteins that are a part of the miRNA induced silencing complex (RISC) and plays an important role in mRNA decay ^54^. It also localizes with AGO and TNRC6 proteins in P-bodies ^55^. L1 ORF1 protein has been previously reported to interact with P-body enriched proteins and RNA ^27,33^. We hypothesize that the absence of AGO2 and mature miRNAs in the miRNA mutant mESCs prevent P-body formation and hinders similarly L1 ORF1 partitioning and LLPS. We think that the observed aggregates in mESCs have evolved as a specialized compartment where diverse activities for L1 RNP metabolism are brought together, which will require further dissection. MOV10 is a 5’ to 3’ RNA helicase ^56^ and its catalytic activity is essential for inhibiting human L1 retrotransposition ^23^. Whether this activity is essential for inducing L1 RNP aggregate formation could provide further mechanistic insight.

Given the plethora of functions MOV10 has been implicated in, it is not surprising that mechanisms have evolved to regulate its expression and activity ^53^. Post-translational modification of MOV10 occurs via ubiquitination in neuron cultures derived from rat hippocampus resulting in its degradation ^57^. Moreover, phosphorylation and acetylation of MOV10 have been observed to occur in human cancer cell lines and speculated to regulate its activity and levels ^53^. Data presented here, to the best of our knowledge, is a first to unveil miRNA mediated post-transcriptional regulation of *Mov10* expression. Since MOV10 expression levels observed in *Dicer*_KO were higher than those in *Drosha*_KO mESCs (Extended Data Fig. 3B) it is possible that expression of MOV10 might also be modulated by microprocessor independent miRNAs. While transient transfection with all four tested miRNAs resulted in downregulation of MOV10, the absence of synergistic effect for miR-16-5p and miR-30-5p may rise from the inherent closeness of the two MREs in the 3’UTR of *Mov10* causing steric hindrance and preventing the large RISC complex from binding the two simultaneously. MREs in *Mov10* for all four tested miRNAs miR-138-5p, miR-30-5p, miR-16-5p and miR-153-3p in mESCs are conserved in the 3’UTR sequence of hMOV10, raising the possibility that this mechanism regulating MOV10 expression may also be conserved in humans. Of note miR-138-5p and miR-153-3p are highly expressed in the human brain ^58^ and both miRNAs are downregulated in brain pathologies from Alzheimer’s Disease patients ^59,60^. Activation of expression and mobility of transposable elements has been reported in a majority of neurological disorders ^61^ and certain cancers ^62^. In case the mode of L1 regulation uncovered here in mESCs is conserved, fine-tuning MOV10 expression in disease conditions using miRNA mimics to downregulate or conversely Antagomirs to upregulate MOV10 expression can afford novel means of therapy.

## Material and Methods

### Cell culture

E14TG2a mESC (ATCC CRL-1821) were used as wild type cells. *Dicer*_KO ^13^ and *Drosha*_KO ^46^ were previously generated from E14TG2a in our laboratory using a paired CRISPR-Cas9 approach ^63^. Cells were cultured in Dulbecco’s Modified Eagle’s Medium (DMEM) (Invitrogen) supplemented with 15% pre-selected batch of FBS (GIBCO) tested for optimal mESCs growth, 1000 U/mL of LIF (Millipore), 0.1 mM of 2-β-mercapto-ethanol (Life Technologies), 0.05 mg/mL of streptomycin, and 50 U/mL of penicillin (Sigma). For routine culturing cells were grown on 0.2% gelatin-coated cell culture grade plastic vessels in the absence of feeder cells. For microscopy coverslips were coated with 10 μg/ml Fibronectin (Sigma, FC010) for at least 2 hours at 37°C, coverslips were washed three times with 1x PBS and cells were seeded 16-18 hours before processing them for microscopy. HEK293T cells were grown in Dulbecco’s Modified Eagle’s Medium (DMEM) (Invitrogen) supplemented with 10% FBS (GIBCO), 0.05 mg/mL of streptomycin, and 50 U/mL of penicillin (Sigma). Concentration of various antibiotics used were as follows 1 μg/ml Puromycin (Sigma), 100 μg/ml Hygromycin (Invitrogen), 250 μg/ml G418 (Sigma), 50 μg/ml Blasticidin (Invitrogen).

### Plasmids

3’UTR sequence of mouse MOV10 transcript ENSMUST00000168015.8 was PCR amplified using Fwd 5’-taggcgatcgctcgaggccacagccgcccgcctt-3’ and Rev 5’-ttgcggccagcggccttttgcatagaacagcattttgt-3’ primers using cDNA generated with random primers from mESCs as template. The PCR product was subcloned into plasmid psiCHECK2 (Promega) previously digested with NotI using the In-Fusion cloning kit (Takara Bio) giving rise to plasmid psiCHECK2-mMov10-3’UTR (addgene 178905). Human MOV10 was PCR amplified with primers Fwd 5’-ggtcggaggcggatccatgcccagtaagttcagctgc-3’ and Rev 5’-gatatctgcagaattctcagagctcattcctccactc-3’ using plasmid pFLAG/HA-MOV10 (addgene 10976) ^64^ as template and subcloned into BamH1 and Xho1 digested pCDNA3-T11-HA plasmid ^65^ a kind gift from Pof. Polymenidou using In-Fusion cloning kit (Takara) to yield plasmid pCDNA3-T11HA-hMOV10-WT (addgene 178907) for transient transfections to over express MOV10 in L1^Up^ and Ctrl cells. Plasmids used for the retrotransposition assay with mneo1 cassette as reporter was pCEP-L1SM (hygro) and with mblast1 cassette was JJ-L1SM WT and JJ-L1SM N21A (hygro), all gifts from Prof. Garcia-Perez.

### Generation of L1^UP^ mESCs using CRISPRa

L1^UP^ mESCs were generated from E14TG2a mESCs using the CRISPRa approach ^66^. Single gRNAs (sgRNAs) were designed using the L1 Tf consensus sequences (Extended Data Fig. 2B) ^67^. Sequence alignments ^6,67,68^ were performed using T-Coffee ^69^. SgRNAs to upregulate L1 Tf were individually sub-cloned into the plasmid pKLV-U6gRNA(BbsI)-PGKpuro2ABFP a gift from Prof. Yusa (addgene 50946), ^70^, using the BbsI restriction site. Guide sequence used for generating Cl1 was 5’-caccgccagagaacctgacagcttc-3’ (addgene nb pending) For Cl2 two guide pairs were used 5’-caccgccagagaacctgacagcttc-3’ (addgene nb pending, same as for Cl1) and 5’-cacccagaggacaggtgcccgccgt-3’ (addgene nb pending). AC95-pmax-dCas9VP160-2A-neo was a gift from Prof. Jaenish (addgene 48227) ^66^. Cells were transfected with 1 μg of each plasmid and 24h hours post transfection they were cultured in presence of puromycin (1 μg/mL) and G418 (250 μg/mL). Single clones were picked one week post transfection. The first screening for selection of L1^UP^ candidates was performed at the protein level for ORF1 expression by immunoblot analysis.

### Ectopic protein expression

L1^UP^ and Ctrl mESC lines were transiently transfected with 2 μg T11HA-hMOV10 plasmid (addgene nb pending) for ectopic expression of hMOV10 or T11HA-EV ^65^ as empty vector control using Lipofectamine 3000 (Invitrogen). Transfection complex was removed 6 hours post transfection. Cells were trypsinized 32 hours post transfection and plated on fibronectin coated cover slips. Samples were processed 48 hours post transfection for Indirect Immunofluorescence (IF).

### MiRNA mimic transfections in mESCs

100,000 *Drosha*_KO mESCs were seeded per well in a 6 well plate in duplicate for respective miRNA mimic transfections. Cells were grown in antibiotic free media and transfected with 20 nM mimic when transfected singly or 10 nM respective mimic for dual transfections using RNAimax reagent (Invitrogen). Cells were harvested 39 hours post transfection and duplicate samples were pooled for protein extraction and subsequent western blot analysis. The following miRNA mimics (Dharmacon, A horizon discovery Group company) were used:

mmu-miR-16-5p 5’-UAGCAGCACGUAAAUAUUGGCG-3’ (C-310511-05-05)
mmu-miR-30e-5p 5’-UGUAAACAUCCUUGACUGGAAG-3’ (C-310466-07-0002)
mmu-miR-138-5p 5’-AGCUGGUGUUGUGAAUCAGGCCG-3’ (C310414-07-0002)
mmu-miR-153-3p 5’-UUGCAUUAGUCACAAAAGUGAUC-3’(C310428-05-0002)
miRIDIAN microRNA negative control 1 (CN-001000-01-05)

### Indirect Immunofluorescence (IF)

Cells grown on coverslips were washed with 1x PBS, fixed with 3.7% formaldehyde (Sigma) in 1x PBS for 10 minutes at room temperature. Post fixation cells were washed three times in 1x PBS and permeabilized with CSK buffer (100 mM NaCl, 300 mM sucrose, 3 mM MgCl_2_, 10 mM PIPES pH 6.8, 0.5% Triton-X) for 4 minutes on ice. After three further washes with 1x PBS, blocking was initiated in 1x PBS supplemented with 1% BSA and 0.1% Tween-20 for 30 minutes at room temperature. Samples were incubated with primary antibody diluted in blocking buffer for 1 hour at room temperature, there after washed three times with 1x PBS-0.1% Tween-20, incubated with secondary antibody diluted in blocking solution for 1 hour and counterstained with 100ng/ml DAPI (Sigma) in 1x PBS for 4 minutes before mounting on slides in Vectashield (Vector labs). The following primary antibodies diluted in blocking buffer were used: rabbit polyclonal anti-ORF1p (1:1000 kind gift from Donal O’Carroll), mouse monoclonal 15C1BB anti-MOV10 (1:500, A500-009A-T Bethyl Laboratories Inc), rabbit polyclonal anti-G3BP1 (1:500 A302-033A, Bethyl Laboratories Inc), rabbit polyclonal anti-LC3B antibody (1:250, 2775, Cell Signaling Technology), rabbit polyclonal anti-DDX6 (1:500, GTX102795, GeneTex), rat monoclonal anti-HA (1:500, 3F10, Roche). Secondary antibody used were Alexa fluor 488 goat anti-rat IgG (1:4000, 11006, life Technologies), Alexa fluor 488 donkey anti-mouse IgG (1:4000, A21202, Life technologies), Alexa fluor 546 donkey anti-rabbit IgG (1:4000, A10040, Life technologies), Alexa fluor 647 donkey anti-mouse IgG (1:4000, A31571, Life technologies). Images were acquired using the Deltavision multiplex system equipped with an Olympus 1X71 (inverse) microscope, pco.edge 5.5 camera and 60x 1.4NA DIC Oil PlanApoN objective. Z stacks were taken 0.2 μm apart, images de-convolved using Softworx software. Further image analysis and processing were performed using ImageJ. Excel (Microsoft) and Prism 9 (Graphpad) were used for data analysis and statistical testing.

### Combined RNA FISH and IF

Cells grown on coverslips were first processed for IF following the protocol described above except all buffers and solution other than the fixative were also supplemented with 10 mM Ribonucleoside Vanadyl Complex (NEB). After incubation with the secondary antibody, cells were fixed with 3.7% formaldehyde in 1x PBS for 10 minutes at room temperature and blocked in 1x PBS supplemented with 1% BSA, 0.1% Tween-20, 2 mM Glycine and 10 mM RVC for 15 minutes. Cells were next washed and incubated in 2x SSC (0.03 M Sodium citrate in 0.3 M Sodium chloride) for 5 minutes. Probe specific for Tf L1 family was labeled with Red-dUTP (Enzo Life sciences) using a nick translation kit (Abbot). 2 μg TFkan plasmid kind gift from Prof. Heard ^71,72^ as incubated with 0.2 mM labelled dUTP, 0.1 mM dTTP, 0.1 mM dNTP mix and 2.5 μl nick translation enzyme in a 50 μl final volume as per guidelines from the kit. The reaction was incubated at 15°C for 15 hours. A PCR purification column (zymogen) was used to clean the probe which was eluted in 50 μl water. The volume of the probe was decreased down to 5 μl using a speed vac, and the probe was diluted in 100 μl hybridization solution (1 part 20x SSC, 2 parts 10 mg/ml BSA, 2 parts 50% Dextran sulfate and 5 parts deionized formamide). The probe solution was denatured at 78°C for 5 minutes, placed on ice for 5 minutes and 7 μl probe was spotted on a pre-baked slide for each sample. During the overnight hybridization at 37°C in a humid chamber the overturned coverslips were sealed using rubber cement. Post hybridization washes were performed with 50% formamide in 2x SSC thrice for five minutes followed by 3 washes with 2x SSC. DNA was counterstained with 100 ng/ml DAPI in 2x SSC and mounted on slides with Vectashield. Image acquisition and analysis was as for IF.

### Western blot analysis

Total cellular protein was extracted from mESC pellets using a NP40 based lysis buffer (1% NP40, 137 mM NaCl, 20 mM Tris-HCl, 1 mM EDTA) complemented with EDTA-free protease inhibitor cocktail (Roche). Protein concentrations were determined by Bradford Assay (Bio-Rad). 10-20 μg of total cellular protein were separated in 8% or 10% SDS-PAGE gels and transferred on PVDF membranes. The following antibodies were used: rabbit polyclonal anti L1 ORF1p (1:5000, gift from Donal O’Carroll), rabbit polyclonal anti-Dicer (1:2000, SAB42000087, Sigma), rabbit polyclonal anti-Argonaute2 (1:2000 C34C6 Cell Signaling Technologies), rabbit anti-Drosha (1:2000, D28B1 Cell Signaling Technology), rat monoclonal anti-HA (1:500, 3F10, Roche), Mouse anti-Tubulin antibody (1:10000, A01410, GenScript), rabbit anti-LaminB1 (1:5000, ab16048, Abcam), rabbit anti-DDX6 (1:2000, GTX102795 GeneTex), anti-rabbit IgG HRP-linked (1:10000 7074, Cell Signaling Technologies), anti-mouse IgG HRP-linked (1:10000, 7076, Cell Signaling Technologies), anti-rat IgG HRP-linked antibody (1:10000, 7077, Cell Signaling Technologies). Immunoblot blot were developed using the Clarify™ Western ECL substrate (BioRad) kit or SuperSignal™ West Femto Maximum Sensitivity Substrate (Thermo Scientific) and detected using ChemiDoc™ MP imaging system (BioRad). All membranes were stained with coomassie to ensure equal loading.

### RT qPCR analysis

Total cellular RNA was extracted from cell pellets using TRizol ® Reagent (Life Technologies). Extract quality was verified by loading 1 μg of total cellular RNA on a 1% Agarose gel. 1 μg cellular RNA was treated with DNase (RQ1 Rnase-Free DNase kit Promega) and reverse-transcribed following the GoScript ™ Reverse Transcriptase Kit (Promega) manufacturer’s instructions. The produced cDNAs were diluted five-fold in distilled water. For each extract, PCR on the *Rrm2* gene were performed, with and without reverse transcriptase treatment, to insure absence of genomic DNA contamination. The quality-controlled cDNAs were diluted two times in distilled water. Amplifications were performed on the Light Cycler ® 480 (Roche) using 2 μL of the diluted cDNAs and the KAPA SYBR ® FAST qPCR Kit Optimized for Light Cycler ® 480 (KAPA biosystems). Differences between samples and controls were calculated based on the 2^−ΔCT^ method. RT-qPCR assays were performed in biological triplicate. Primers utilized for the RT-qPCR assays are as follows: Rrm2fwd 5’-ccgagctggaaagtaaagcg-3’, Rrm2rev 5’-atgggaaagacaacgaagcg-3’, Mov10fwd 5’-gacgatttacaaccacgacttca-3’, Mov10rev 5’-gccagatttgcgatcttcattcc-3’, Dicerfwd 5’-ccgatgatgcagcctctaatag-3’ Dicerrev 5’-tccatctcgagcaattctctca-3’, L1-Tffwd 5’-cagcggtcgccatcttg-3’, L1-Tfrev 5’-caccctctcacctgttcagactaa-3’, L1-Afwd 5’-ggattccacacgtgatcctaa-3’, L1-Arev 5’-tcctctatgagcagacctgga-3’, L1-Gffwd 5’-ctccttggctccgggact-3’, L1-Gfrev 5’-caggaaggtggccggttgt-3’, L1-ORF1fwd 5’-actcaaagcgaggcaacact-3’ L1-ORF1rev 5’-ctttgattgttgtgccgatg-3’, L1-ORF2fwd 5’-ggagggacatttcattctcatca-3’, L1-ORF2rev 5’-gctgctcttgtatttggagcataga-3’.

### Northern Blot analysis

Northern blot analysis was performed as previously described ^13,73^. 30 μg of total RNAs extracted using Trizol were run on a denaturing 1% Agarose gel with 1% Formaldehyde. Following capillary transfer to nylon membranes overnight the membrane was cross-linked by UV radiation. PerfectHybTM Plus was used for pre hybridization blocking and hybridization at 42°C. Post hybridization washes were performed in 2x SSC + 0.1% SDS. For detection of full-length L1 transcripts, random primer extension labeling was carried out. DNA used for the reaction was PCR amplified using E14TG2a mESCs genomic DNA as template and L1specifc primers Fwd 5’-gagtttttgagtctgtatcc-3’ and Rev 5’-ctctccttagtttcagtgg-3’.

### Dual luciferase reporter assay

70,000 HEK293T cells were plated per well in a 24 well plate 16 hours prior to transfection with Lipofectamine 2000 (Invitrogen). 0.5 μg of plasmid psiCHECK2-3’UTR-WT-Mov10’UTR was co-transfected with 50 nM indicated miRNA mimics or control mimic. Transfection complexes were removed 6 hours post transfection. Luciferase activity was measured on a GloMax® Discover Multimode Microplate Reader (Promega, USA) after processing cells using the Dual-Glow Luciferase Assay kit (E2920 Promega, USA) 48 hours post transfection. Results are means and error bars are standard deviation (SD) from three to four independent experiments.

### Retrotransposition reporter and colony forming assays

1×10^6^ L1UP and Ctrl mESCs were seeded in 10 cm dish 16 hours prior to transfection with 6 μg of JJ-L1SM (WT and L1N21A) plasmid using Lipofectamine 3000 (Invitrogen). Media exchange was initiated 6 hours post transfection and hygromycin supplemented media was added 48 hours post transfection to select for stably transfected cells. Once the mock transfected cells were dead, 150,000 hygromycin resistant cells were seeded per well in a 6 well plate in triplicate and grown in media *sans* hygromycin for 16 hours after which the media was supplemented with Blasticidin. Media exchange with fresh antibiotics was performed every 48 hours for approximately 15 days, when individual Blasticidin resistant colonies were visible with the naked eye. Cells were washed with 1x PBS and stained with 1% crystal violet blue, 1% formaldehyde, 1% methanol for 20 minutes at room temperature, followed by washes with tap water. Plates were air dried and imaged using the ChemiDocTM MP system (BioRad). Individual colonies were counted using ImageJ. Results are means and error bars are SD from three independent transfections.

Transient transfections of reporter plasmids were carried out using Lipofectamine 3000 (Invitrogen) when co-transfections with miRNA mimics or plasmids for ectopic expression of hMOV10 were assayed for retrotransposition. 500,000 cells were seeded for transient transfection with 6 μg of reporter plasmid and either 10 nM mimic for mmu-miR-16-5p + 10 nM mmu-miR-153-3p mimic or 6 μg of plasmid T11HA-EV or T11HA-hMOV10. Media exchange was initiated 6 hours post transfection. 39 hours post transfection cells were grown in media supplemented with antibiotic resistance encoded by the respective cassette. Subsequent media exchanges, staining and counting of colonies, was the same as stated for stably transfected cells. Results are means and error bars are SD from three independent transfections.

## Acknowledgments

We would like to thank the members of the Ciaudo lab and Dr Tobias Beyer for fruitful discussions and the critical reading of this manuscript. This work was supported by the Swiss National Science Foundation (grants 31003A_173120 and 310030_196861) and Novartis Foundation for medical-biological Research (grant 19A018) to C.C. In addition, R.A and C.C. were supported by the NCCR RNA and Disease. We also want to thank the ScopeM facility at ETH Zürich for their support with microscopy and image analysis.

## Author Contributions

Conceptualization, RA, MB and CC, laboratory experiments RA, MB, LP, CH; writing original draft preparation, RA and CC; writing, review and editing, CC; visualization, RA, MB and CC; supervision, CC; funding acquisition, CC. All authors have read and agreed to the published version of the manuscript.

---

**Extended Data Fig. 1.**
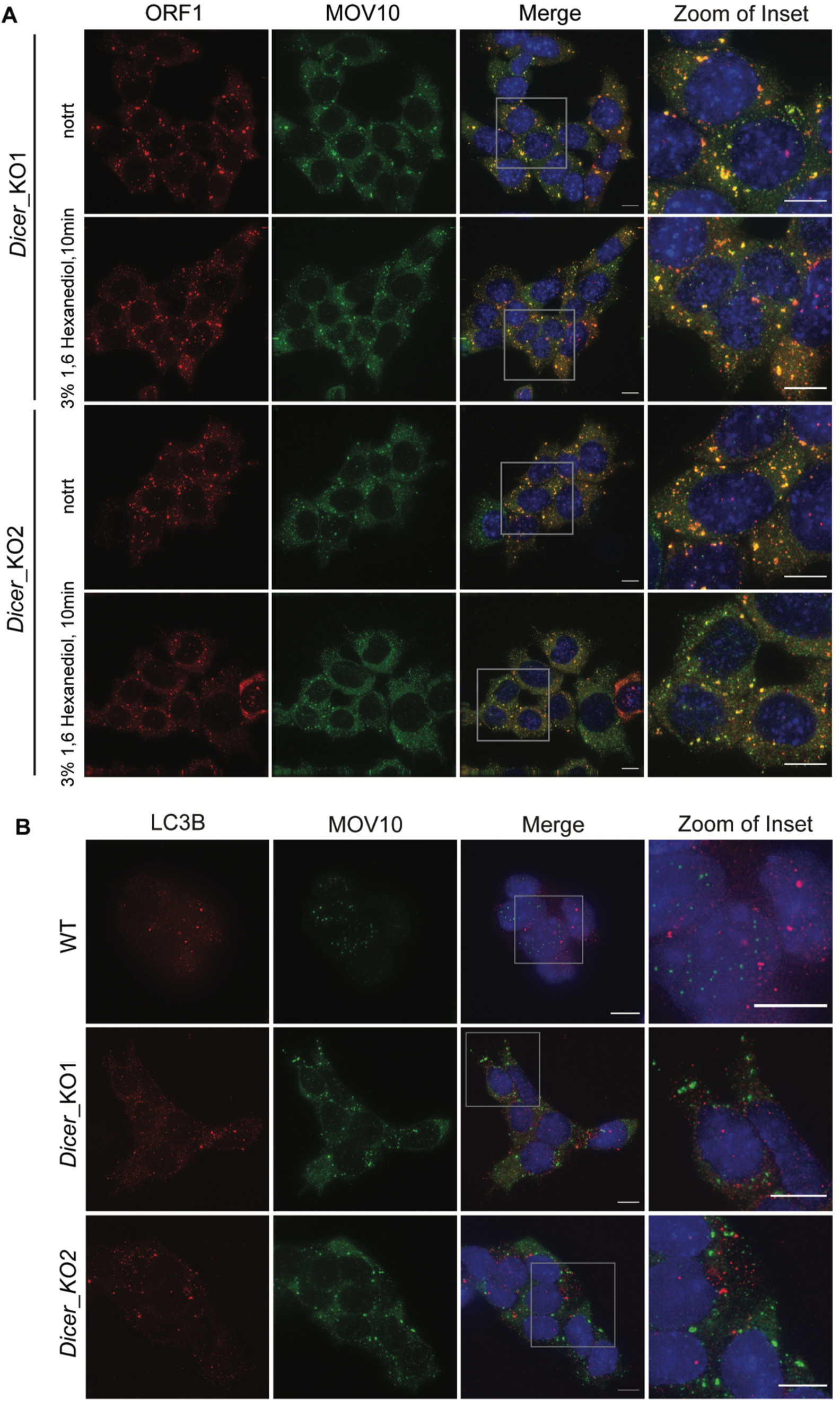
L1 RNP cytosolic aggregates are not sensitive to treatment with 3% 1,6 Hexanediol. **(A)** WT and *Dicer*_KO mESCs were treated the 3% 1,6 Hexanediol for 10 minutes to assess ability of L1 RNP to phase separate. Maximum intensity projections across Z stacks of example images from indicated mESCs immunostained for L1 ORF1 (red) and MOV10 (green) with nuclei stained with DAPI (blue). The lack of any discernible change in L1 ORF1-MOV10 foci formation indicates absence of LLPS for L1 ORF1-MOV10 foci. Images are representative of 3 independent experiments. Grey box mark position of the insets. **(B)** Maximum intensity projections across Z stacks of example images from indicated mESCs immunostained for LC3B (red) and MOV10 (green) with nuclei stained with DAPI (blue). The absence of any co-localization of LC3B with MOV10 in the tested cell lines indicate that the L1 RNP foci are not autophagosomes. The grey square depicts position of inset. Images are representative of 3 independent experiments. Scale bar 5 μm.

**Extended Data Fig. 2.**
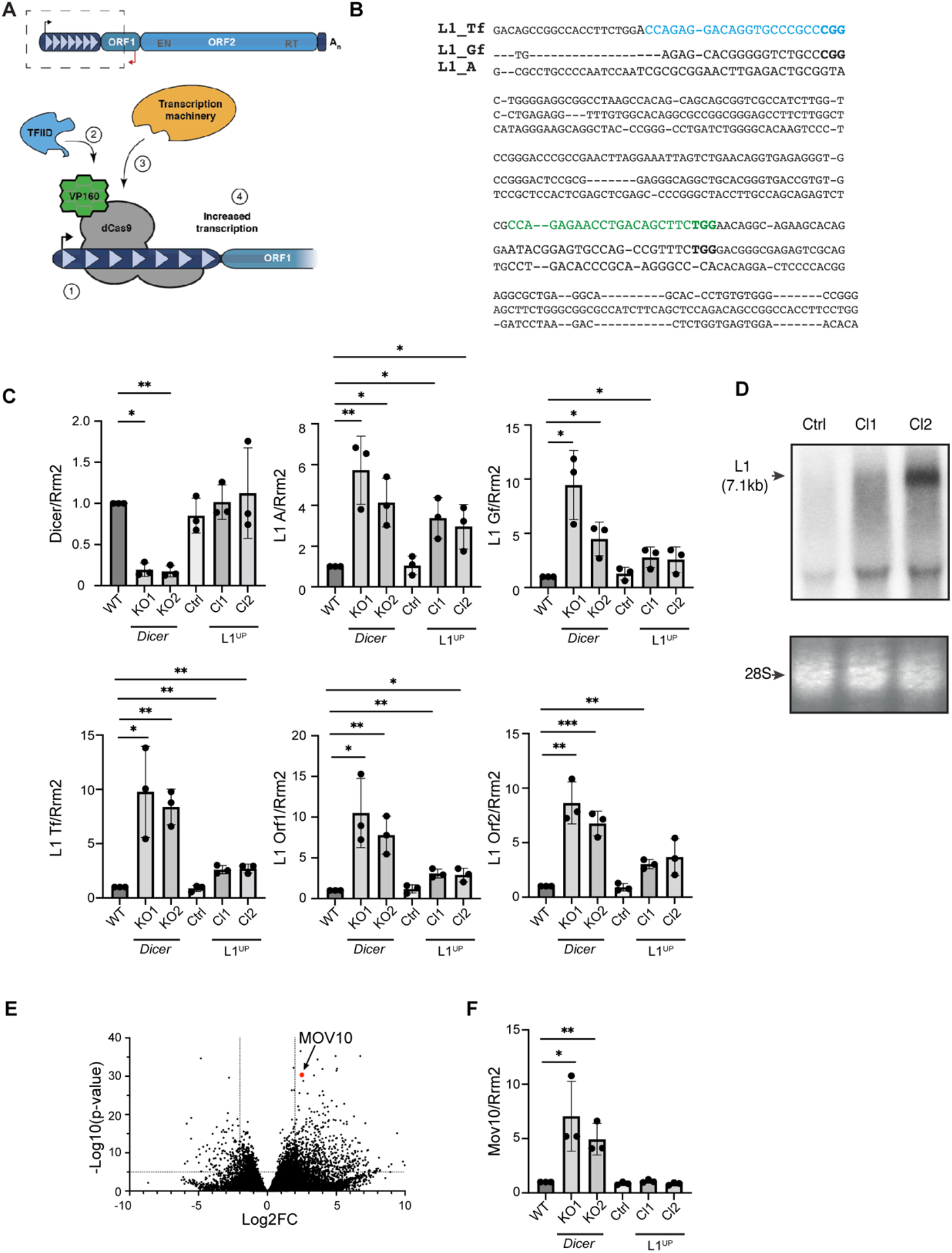
Generation of mESCs upregulating L1 expression using CRISPRa. **(A)** Schematic depicting full length L1 element and summary of CRISPRa. To generate L1^UP^ cells, mESCs were co-transfected with plasmid encoding catalytically dead Cas9 protein (dCas9) fused to VP160 and sgRNAs that (1) targeted the fusion protein to the 5’UTR sequence of Tf L1 family allowing (2) recruitment of transcription factors and (3) transcription machinery to (4) upregulate L1 transcription. **(B)** Sequence alignment of 5’UTR sequences of murine L1 Tf, Gf and A subfamilies. The two sgRNA sequences used to upregulate L1 expression are indicated in blue and green, with protospacer adjacent motifs (PAM) in bold. **(C)** RT qPCR analysis to quantitate *Dicer*, and L1 RNA expression levels in the depicted cell lines. *Rrm2* was utilized for normalization, and graphs depict fold change in transcript levels in the indicated cell lines as compared to WT which was set to one. Bar graphs show means from 3 independent experiments, error bars are standard deviations, p-values were computed using unpaired t-test comparing results from individual cells to WT mESCs. Asterisk are p-values * < 0.05, ** < 0.001, *** <0.0005. **(D)** Northern blot analysis probed for L1 RNA to assess L1 transcript length and expression levels in the engineered L1^UP^ Cl1, Cl2 as compared to Ctrl cells. Arrow points to full length L1 transcript. Ethidium bromide staining of 28S RNA was used to confirm equal loading. **(E)** Differential gene expression from RNA-seq analysis of *Dicer*_KO vs. WT mESCs plotted using previously published data ^13^. Each dot represents a single gene. Position for *Mov10* in the graph is marked. Values for Log2 fold change (Log2FC) were plotted on the x-axis and Log10 of the p-value on the y-axis. **(F)** RT qPCR analysis to confirm upregulation of *Mov10* in *Dicer*_KO cells. No change in *Mov10* transcript levels were observed in Ctrl, L1^UP^ Cl1, Cl2 as compared to WT mESCs. *Rrm2* was utilized for normalization, and graphs depict fold change in transcript levels in the indicated cell lines as compared to WT which was set to one. Bar graphs show means from 3 independent experiments, error bars are standard deviations, p-values were computed using unpaired t-test comparing results from individual cells to WT mESCs. Asterisk are p-values * < 0.05, ** < 0.001.

**Extended Data Fig. 3.**
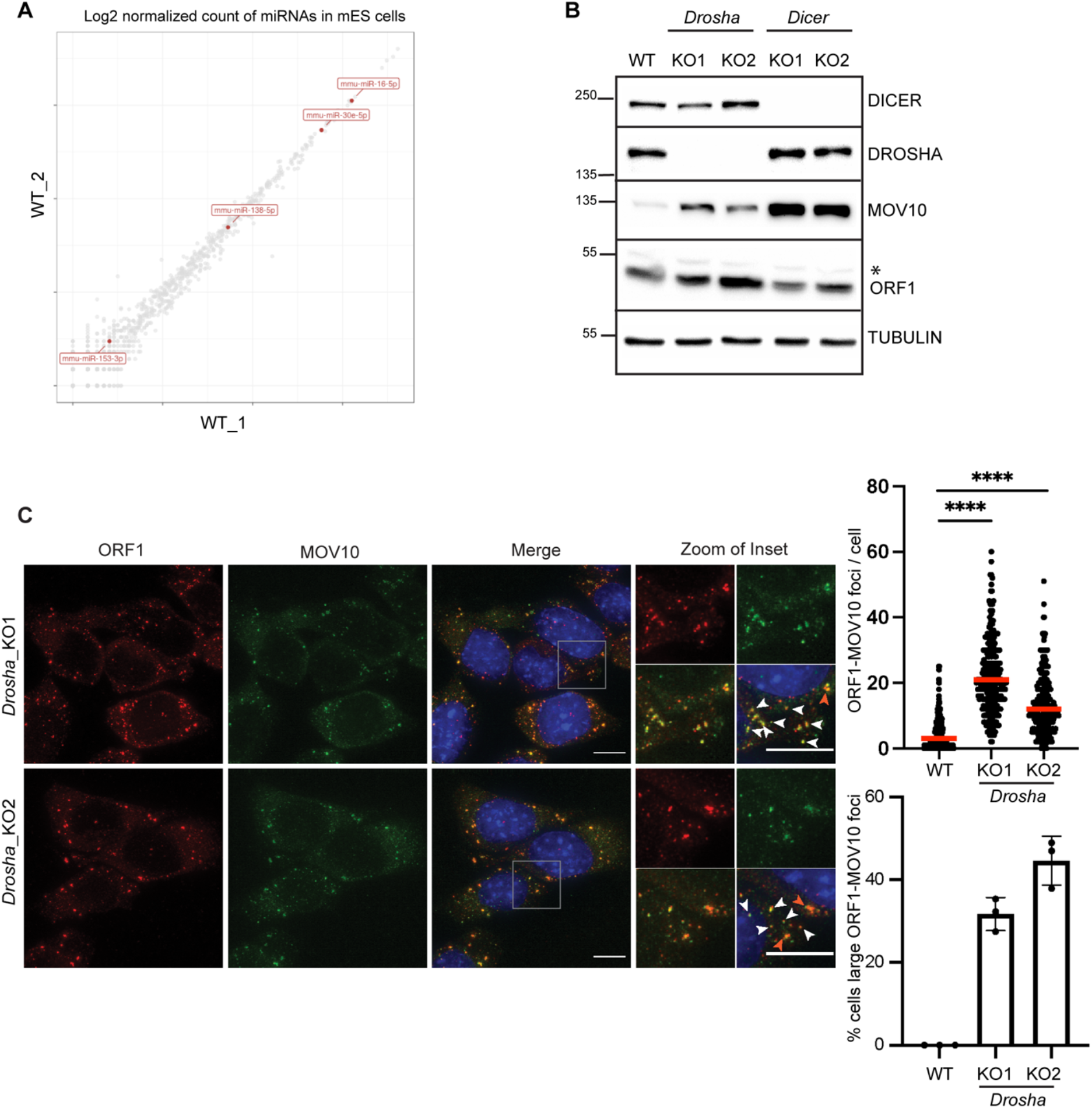
Upregulation of L1 ORF1 and MOV10 in the absence of miRNAs in *Drosha*_KO mESCs induces L1 RNP aggregation in the cytoplasm. **(A)** Log2 normalized count of miRNAs in WT mESCs from previously published small RNA-seq data ^44^. Each dot depicts a single miRNA and miRNAs predicted using TargetScan ^43^ to regulate *Mov10* expression are shown in red. **(B)**Western Blot analysis to assess expression of L1 ORF1 and MOV10 in the indicated cell lines, immunoblot with TUBULIN served to control for loading. Membranes were probed with anti-DICER and anti-DROSHA antibodies to confirm the deletion status of the cells. Upregulation of L1 ORF1 and of MOV10 was observed in *Drosha*_KO relative to WT mESCs. Asterisk marks non-specific band recognized by ORF1 antibody **(C)** Maximum intensity projections across Z stacks of example images from *Drosha*_KO mESCs immunostained for L1 ORF1 (red), MOV10 (green) and nuclei stained with DAPI (blue). White arrow heads point to cytoplasmic foci where L1 ORF1 and MOV10 co-localize. Red arrow heads point to relatively larger sized L1 RNP foci. Data collected from 288 *Drosha*_KO1, 299 *Drosha*_KO2 cells from three independent experiments are depicted as scatter plots where circles are single data points representing number of co-localized L1 ORF1-MOV10 foci in the cytoplasm per cell, red bar is median for the distribution. Data for WT cells for comparison is the same as in Figure 1B. P-value was determined using Mann-Whitney *U* test and **** represent p-value < 0.0001. Bar graphs are mean values of percentage of cells with large L1 ORF1-MOV10 foci co-localizing in the cytoplasm. Dots represent data from three independent experiments, error bars are standard deviations. Scale bar 5 μm.

